# EGFR activation in cholangiocytes promotes extrahepatic bile duct regeneration after injury

**DOI:** 10.1101/2025.04.22.649788

**Authors:** Ashley N. Calder, Takuki Sakaguchi, Mirabelle Peter, John Tobias, Timothy Frankel, Nataliya Razumilava

## Abstract

**Background & Aims:** The epidermal growth factor (EGF) receptor family of tyrosine kinases regulates development and homeostasis of digestive organs including the liver and bile ducts. It consists of four receptors, EGF receptor (EGFR) and erythroblastoma oncogene B 2-4 (ERBB2–4), and their corresponding ligands. EGF signaling promotes intrahepatic cholangiocyte proliferation, bipotent cell transdifferentiation into cholangiocytes, bile duct branching, and cholangiocarcinoma (CCA) aggressiveness.

The EGF family signaling contribution to extrahepatic bile duct (EHBD) regeneration is not well defined. This work is aimed at determining the fundamental role of the EGF signaling network in the biliary proliferative response to EHBD obstruction.

**Approach:** We used mouse bile duct ligation to model obstructive EHBD injury, and human and mouse EHBD organoids for *in vitro* studies. We tested activating and inhibitory paradigms with recombinant EGF family ligands and receptor antagonists.

Transcriptomic and immunohistochemistry analyses informed EGF signaling changes and cellular localization at homeostasis and after obstruction.

**Results:** At homeostasis, the EHBD expressed EGFR ligands *Tgfa, Btc, Hb-egf* and *Nrg4* in cholangiocytes, and *Egf* in stromal cells. *Erbb2* and *Erbb3* were predominant receptors expressed in cholangiocytes and *Egfr* in stromal cells at baseline. After EHBD obstruction, injury-induced biliary hyperproliferation was associated with increased abundance of *Areg*, *Hb-egf*, *Tgfα* and *Btc* ligands and *Egfr* receptor in cholangiocytes with resulting epithelial EGFR activation. In biliary organoids, EGFR ligands induced organoid growth, and inhibition of EGFR, but not ERBB2, dampened cholangiocyte proliferation. Accordingly, EGFR inhibition in mice led to a decrease in the biliary proliferative response after EHBD obstruction.

**Conclusion:** The obstruction-induced biliary proliferation is an EGFR-mediated response suggesting context-and receptor-specific EGF signaling network involvement in EHBD regeneration after injury.

## INTRODUCTION

The biliary tree is the residence of a wide range of pathologies collectively named cholangiopathies. Cholangiopathies are associated with biliary obstruction, inflammation, biliary hyperproliferation and extracellular matrix remodeling, which can progress to liver fibrosis and failure requiring liver transplant or to biliary cancer (1). It is of major importance to develop strategies supporting biliary repair after injury to prevent cholangiopathy development and progression.

Organ regeneration aims to restore the organ function after injury through recovery of its cellular and structural integrity. Cell proliferation is a common regenerative response to insults. In the hepatobiliary system, liver regeneration is the most studied, in part owing to the liver’s striking ability to recover its pre-resection hepatocyte volume in just three weeks (2–4). Cholangiocytes, similar to hepatocytes, are mitotically quiescent at homeostasis, yet they exhibit a prominent proliferative response to obstructive injury, as we and others had shown using a mouse bile duct ligation (BDL) model (5–7).

Surprisingly, the mechanisms regulating regeneration and biliary epithelium proliferation in large bile ducts, including the extrahepatic bile duct (EHBD), are less studied. Large perihilar and distal bile ducts compose the EHBD. There are pathological differences in diseases of intra-and extrahepatic bile ducts due to their distinct anatomic location, histological composition, and embryonic origin (8). This work aims to uncover fundamental mechanisms supporting EHBD repair through the proliferative response to injury.

Developmental signaling pathways, including WNT, Hedgehog and Hippo signaling, are implicated in large bile duct responses to injury (9, 10). Our group has recently reported tightly regulated intercellular communication supporting EHBD proliferation after obstruction (5). In this cellular crosstalk, Hedgehog signaling originates from injured cholangiocytes to communicate to receptive fibroblasts recruiting neutrophils into the damaged bile duct to promote cholangiocyte proliferation. Cholangiocyte autocrine signaling through upregulation of WNT ligands with resulting β-catenin transcriptional programming activation can also support biliary proliferation (6). While both Hedgehog and WNT pathways play a role in bile duct regeneration, their inhibition does not completely abrogate the proliferative response of cholangiocytes suggesting involvement of other mechanisms.

In the past, numerous studies have explored the epidermal growth factor (EGF) signaling network in liver biology. EGF signaling consists of four receptor tyrosine kinases, EGF receptor (EGFR) and erythroblastoma oncogene B (ERBB) 2-4 (11), also referred as human epidermal growth factor receptors (HER1-4) in humans. EGF is a prototypic EGFR signaling ligand. Transforming growth factor-α (TGFα), betacellulin (BTC), heparin-binding EGF-like growth factor (HB-EGF), amphiregulin (AREG) and epiregulin (EREG) ligands have binding specificity for EGFR (TGFA, BTC, HB-EGF, AREG, and EREG) and ERBB4 receptors (BTC and HB-EGF) (12). Notably, ERBB2 lacks a ligand-binding domain on its extracellular region and, therefore, requires heterodimerization with other EGF network receptors. Neuregulin 1 and 2 (NRG1,2) are ligands for ERBB3 and NRG1-4 for ERBB4. (12, 13). In the liver, EGF was shown to promote hepatoblast differentiation into cholangiocytes, branching in tissue-engineered bile ducts, and support biliary organoid cultures (14). EGFR and AREG were shown to be increased in livers of patients with primary biliary cholangitis and primary sclerosing cholangitis associated with biliary hyperproliferation and protect from cholestatic liver injury in mice by supporting cholangiocyte proliferation (15). Genetic inhibition of EGFR in partial hepatectomy dampened cellular proliferation by inhibiting the cell cycle and delayed hepatocellular carcinoma development in a chemical mouse model (16). Lastly, a recent study demonstrated that the primary cilia defect in cholangiocytes decreases EGFR turnover with resulting sustained EGFR signaling and aberrant biliary hyperproliferation observed in polycystic liver disease and intrahepatic cholangiocarcinoma (CCA) (17). These studies suggest the role of EGF signaling network in liver responses to injury and in liver cancer. Despite the body of research on EGF signaling network in the liver and intrahepatic bile ducts, its status, cellular localization and effects in EHBD injury are not well described. Accordingly, we hypothesized that the EGF signaling is upregulated in cholangiocytes during obstructive EHBD injury to promote bile duct regeneration with biliary proliferation.

In the present study, we used BDL to model obstructive EHBD injury as other currently available models of cholestatic liver injury are not known to recapitulate the proliferative phenotype in EHBDs (18, 19). We used human and mouse EHBD organoid models to assess the direct effects of EGF family signaling activators and inhibitors on cholangiocytes. We report activation of the EGF signaling network in obstructed EHBDs with increased abundance of EGFR ligands, and phosphorylated EGFR in cholangiocytes. EGFR ligands exhibited a redundant role in promotion of EHBD organoid proliferation. Through testing of an inhibitory paradigm, we determined dependence of the biliary proliferative response on EGFR, and not ERBB2, activation suggesting receptor-and context-specific effects. Together, our study unveiled the landscape of EGF family signaling components in the EHBDs and showed the prominent role of activated EGFR in promoting EHBD cholangiocyte proliferation in response to biliary obstructive injury.

## MATERIALS AND METHODS

### Mice

All mouse experiments were approved by the University of Michigan’s Institutional Animal Care and Use Committee. All mice were bred on a C57BL/6 background and housed in a specific pathogen-free environment with a 12:12-hour light-dark cycle in ventilated caging with enviro-dri absorbent or cotton squares as enrichment and provided ad libitum access to food (Purina LabDiet 5L0D; St. Louis, MO) and water. All mice were 2-6 months old at the time of experiment, both male and female mice were used and sex and age matched, when possible. Animals were euthanized according to institutional guidelines. Acute obstructive EHBD injury was induced by BDL as was previously described (5). For pharmacologic inhibition of EGFR signaling, EGFR inhibitor erlotinib (LC Laboratories, Cat#: E-4007, Woburn, Massachusetts) was resuspended at 20 mg/mL in PEG400 immediately before the experiment and administered by gavage at 100 mg/kg starting 24 hours prior to the BDL and every 24 hours thereafter. To assess cellular proliferation, 5-ethynyl-2’-deoxyuridine (EdU; Lumiprobe, Cat#: 10540, Cockeysville, MD) was prepared according to the manufacturer’s instructions and administered to mice intraperitoneally 2 hours prior to euthanasia as previously described (6).

### Human and mouse organoid cultures

EHBD biliary organoid cultures were generated and maintained as previously described with minimal modifications (6). In brief, mouse and human EHBDs were minced and dissociated using the dissociation media Liberase TL (Millipore Sigma, Cat#: 5401020001, Burlington, MA) and grown in organoid growth media containing 10% L-WRN and 30% L-WRN conditioned media for mouse and human organoids respectively, and 50ug/mL EGF for human organoid maintenance. For experiments with EGF signaling modulators, human and mouse organoids were cultured in EGF-free, serum-free, and conditioned-free medium supplemented with 3 µM CHIR-99021 to maintain WNT signaling (Apex Bio, Cat#: A3011, Houston, TX) as indicated in the experimental timelines. EGFR ligands TGFα (100-16A), BTC (100–50), HB-EGF (100–47), AREG (100-55B), EREG (100–04), NRG1 (100–03) were purchased from Peprotech (Cranbury, NJ) and resuspended per the manufacturer’s protocol in 0.1% bovine serum albumin and stored at-20°C. Ligands or vehicle control (0.1% bovine serum albumin) were diluted on the day of the experiment and added to the culture medium. The initial ligand doses were determined based on the published literature (20–22). For pharmacologic inhibition of EGFR, erlotinib (LC Laborotories, E-4007; Woburn, MA) was resuspended in dimethyl sulfoxide at a concentration of 10 mM and stored at-20°C. Media containing erlotinib or vehicle control (dimethyl sulfoxide) were made and added to the culture medium 24 hours after organoid seeding, concentrations were determined based on published literature (22). Trastuzumab, an ERBB2 inhibitor, was reconstituted in sterile phosphate-buffered saline to a concentration of 10mg/mL and stored at 4°C. Medium containing trastzumab or control (Human IgG isotype) was made and added to culture media 24 hours after organoid seeding, concentrations were chosen based on published literature (23)). Organoid and cell culture growth was assessed with CellTiter-Glo 3D Viability Assay (Promega, G9681; Madison, WI) and the CellTiter-Glo 2.0 (Promega, G924A) Viability Assay respectively through measurement of Adenosine triphosphate (ATP) levels as previously described (6). Organoid number and growth rates were accessed by enumeration of organoids with ImageJ (NIH) using images from at least 3 wells taken with a stereomicroscope (Olympus, SZX16) and digital microscope camera (Olympus, DP72) using cellSens software (Olympus, Center Valley, PA).

### BT474 breast cancer line cell culture

The immortalized ERBB2 positive breast cancer cell line BT474 (23, 24) was a kind gift of Dr. Carole Parent (University of Michigan). Cells were maintained in RPMI culture medium containing 10% FBS and 1% Pen-strep and seeded in a 48-well plate at 30,000 cells/well. Media containing 10 ug/mL trastuzumab (MedChemExpress, HY-P9907; Monmouth Junction, NJ) (23) or an human IgG isotype (Invitrogen, 027102) control was added to cells 24 hours after seeding and every two days until the final analysis.

### Immunofluorescent analysis

Mouse EHBDs were isolated 24 post-BDL, washed in phosphate-buffered saline, fixed in 10% neutral buffered formalin (Sigma-Aldrich, HT5012) at 4°C overnight and processed for immunofluorescence analysis as previously described (25). Primary and secondary antibodies used for immunofluorescence analysis are listed in Suppl. Table 1. Hematoxylin and eosin (H& E; Vector Labs, H-3502; Newark, CA) staining was done following manufacturer’s protocol (Vector Labs). For DAB staining Vectastain Elite ABC kit, Peroxidase (Vector Labs, PK6100) and DAB substrate kit (Vector Labs, SK4100) were used according to the manufacturer’s instructions. Biliary proliferation was assessed by labeling proliferating cells with EdU and using a Click-it Kit (Invitrogen, C10337) according to the manufacturer’s protocol. Epithelial cell compartment was determined by positive cytokeratin 19 (KRT19) staining. Epithelial and stromal cell compartments were analyzed for the percentage of EdU-incorporating cells as a proportion of 4’,6’-diamidino-2-phenylindole (DAPI)-marked cells in each compartment.

### Transcriptomic analyses

EHBD organoid RNA was extracted as previously described using the Rneasy mini kit (Qiagen, 74106; Germantown, Maryland) with a DNase treatment (Qiagen, 79254; Germantown, MD) according to manufacturer’s instructions (6). The quantitative reverse-transcriptase polymerase chain reaction (qRT-PCR) analysis of mouse EHBD organoids was performed using iScript cDNA Synthesis kit (Bio-Rad, 1708891; Hercules, CA) for cDNA generation and iTaq Universal SYBR Green Supermix (Bio-Rad, 1725121) for mRNA abundance examination. Primer sequences were designed using the NCBI Primer Blast tool (National Library of Medicine, Bethesda, MD) (Suppl. Table 2) and prepared by Integrated DNA Technologies (Coralville, IA). Expression data were normalized to *18s* ribosomal RNA. All biological samples were analyzed in technical triplicates. For the transcriptomic analysis of the whole mouse EHBD by bulk RNA sequencing (RNA-seq) and single cell RNA-seq (scRNA-seq), we conducted a secondary analysis of our published datasets GSE280724 and GSE280889 (6). To evaluate mRNA abundance of EGF family signaling components in human organoids, we mined the published dataset E-MTAB-7569 (26). Analyses of datasets were performed as previously described (6). Dot plots of scRNA-seq datasets were performed in R-studio version 2023.06.1 with the Seurat package version 4.1.1. to visualize gene transcripts within cell populations.

### Gene Set Enrichment Analysis (GSEA)

GSEA (v4.3.3) was run in preranked mode with the DESeq2 statistic as the ranking metric for the BDL vs sham contrast in the bulk RNAseq dataset. Pathways from the hallmark and the c2.cp.biocarta.v2024.1.Hs.symbols collections were scored using 1000 permutations (27). Results for pathways of interest were presented individually.

### Statistical analysis

Unpaired Student *t*-test or one-way analysis of variance (ANOVA) were used to determine statistical significance. Values are presented as mean ± standard deviation (SD) and *P*-value significance was set at *P*<0.05. Control groups are represented by either vehicle-treated samples or sham procedures. All experiments included at least 3 mice per group for all analyses and 3 independent organoid culture lines except for discovery of the activating or inhibiting dose of the pharmacologic agent where at least 3 technical replicates were used. All statistical analyses were performed using Prism v.10 (GraphPad). Significant pathway enrichment was determined using the FDR q-value generated from the GSEA software (27).

## RESULTS

### EGF signaling network is upregulated in obstructed EHBDs

To determine the potential role of the EGF family signaling network in EHBD regeneration, we employed the mouse model of biliary obstruction, BDL, and examined EHBDs at baseline (sham) and post-BDL using histologic and transcriptomic analyses 24 hours post-surgery (**Fig. 1A**). We previously reported that the peak mouse EHBD proliferative response occurs at the 24-hour timepoint (5, 6). Accordingly, we demonstrated epithelial compartment expansion through an increase in the cholangiocyte marker, KRT19, 24 hours after BDL using immunohistochemistry and bulk RNA-seq analyses (**Fig. 1B**).

**Fig. 1.**
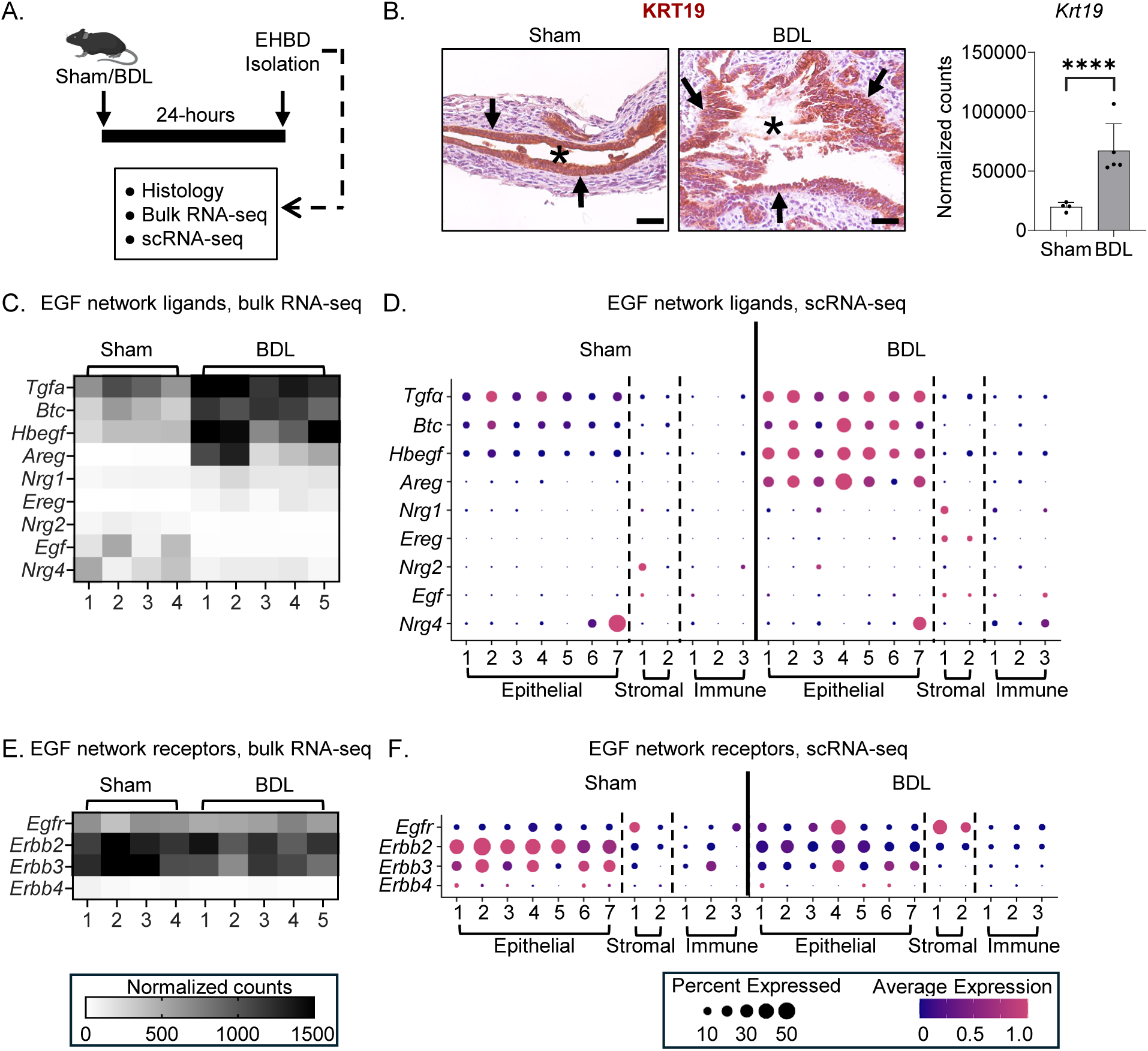
EHBD obstruction results in changes in EGF signaling ligand and receptor expression. Experimental design for wild type (WT) mice that underwent 24-hour sham or bile duct ligation (BDL) (A). Extrahepatic bile ducts (EHBDs) examined for the epithelial marker keratin 19 (KRT19) with immunohistochemistry (brown; B, left) and bulk RNA-seq *Krt19* gene expression (B, right). Epidermal growth factor (EFG) signaling ligand (C,D) and receptor (E,F) expression from bulk RNA-seq (left) or scRNA-seq (right) from EHBDs 24 hours post-surgery. Asterisks mark EHBD lumen. n = 3-5 mice/group. The data are presented as the mean ± SD. \*\*\*\**P* < 0.0001. Arrows mark KRT19+ cholangiocytes. Scale bars, 50μm.

To understand the status of EFG signaling network in homeostatic and injured EHBDs, we analyzed transcriptional expression and cellular localization of ligands with bulk RNA-seq and scRNA-seq. We demonstrated a significant increase in the abundance of *Tgfα, Btc, Hbegf,* and *Areg* ligand-encoding genes post-BDL with bulk RNA-seq (**Fig. 1C**). scRNA-seq analysis mapped these ligands primarily to epithelial cell clusters at homeostasis and after injury (**Fig. 1D**). While *Nrg1* and *Ereg* were expressed at low levels at homeostasis, mapped to fibroblasts and had a statistically significant increase, their mRNA abundance post-BDL remained low relative to other ligands (**Fig. 1C, D)**.

Interestingly, *Egf* and *Nrg2* mapped mainly to fibroblasts and *Nrg4* mapped predominantly to epithelial cells and were expressed at low levels at homeostasis, however their abundance decreased after injury (**Fig. 1C, D**). Thus, these findings suggested an association between biliary proliferation (**Fig. 1B**) and an upregulation of EGFR and ERBB4 receptor ligands in injured EHBDs.

Transcriptomic analysis of our bulk and scRNA-seq datasets before and after BDL showed that receptor-encoding genes *Egfr*, *Erbb2* and *Erbb3* are expressed at high levels in the mouse EHBD at homeostasis (**Fig. 1E**). In cholangiocytes, *Erbb2* and *Erbb3* were the predominant receptors expressed at homeostasis, but their biliary abundance decreased in injured EHBDs. Importantly, *Egfr* abundance increased in cholangiocytes post-BDL suggesting differences in EGFR localization in the EHBD at homeostasis and after damage (**Fig. 1G**). *Erbb4* expression was almost absent in mouse EHBDs (**Fig. 1E**). These data suggested an upregulation of genes encoding EGFR and its ligands in cholangiocytes in the EHBD injury response.

We hypothesized that a surge in ligand expression promotes EGFR activation. As mouse EHBD tissue size and composition with a minimal proportion of epithelial cells prevents effective western blot analysis, we examined sham and BDL mouse EHBDs for phosphorylated (activated) EGFR (pEGFR) by immunostaining. While there was minimal to none phosphorylated EGFR at homeostasis, we observed high expression of phosphorylated EGFR with primary localization to cholangiocytes post-BDL (**Fig. 2A**). To further test our hypothesis that EGFR is activated post-BDL, we conducted a gene set enrichment analysis (GSEA) of the bulk RNA-seq datasets from sham/BDL EHBDs with a focus on PI3K/AKT/MTOR and MAP kinase signaling (**Fig. 2B, C; Suppl. Fig. 1**), which are downstream from EGFR (28). As anticipated, we observed significant enrichment in PI3K/AKT/MTOR signaling post-BDL as compared to sham EHBDs (**Fig. 2B, C**). While there was no statistically significant enrichment in the MAP kinase pathway post-BDL (**Suppl. Fig. 1**). Together, these data supported our hypothesis that injury-induced upregulation of EGFR ligands results in activation of EGFR in cholangiocytes with an increase in PI3K/AKT/MTOR signaling post-BDL.

**Fig. 2.**
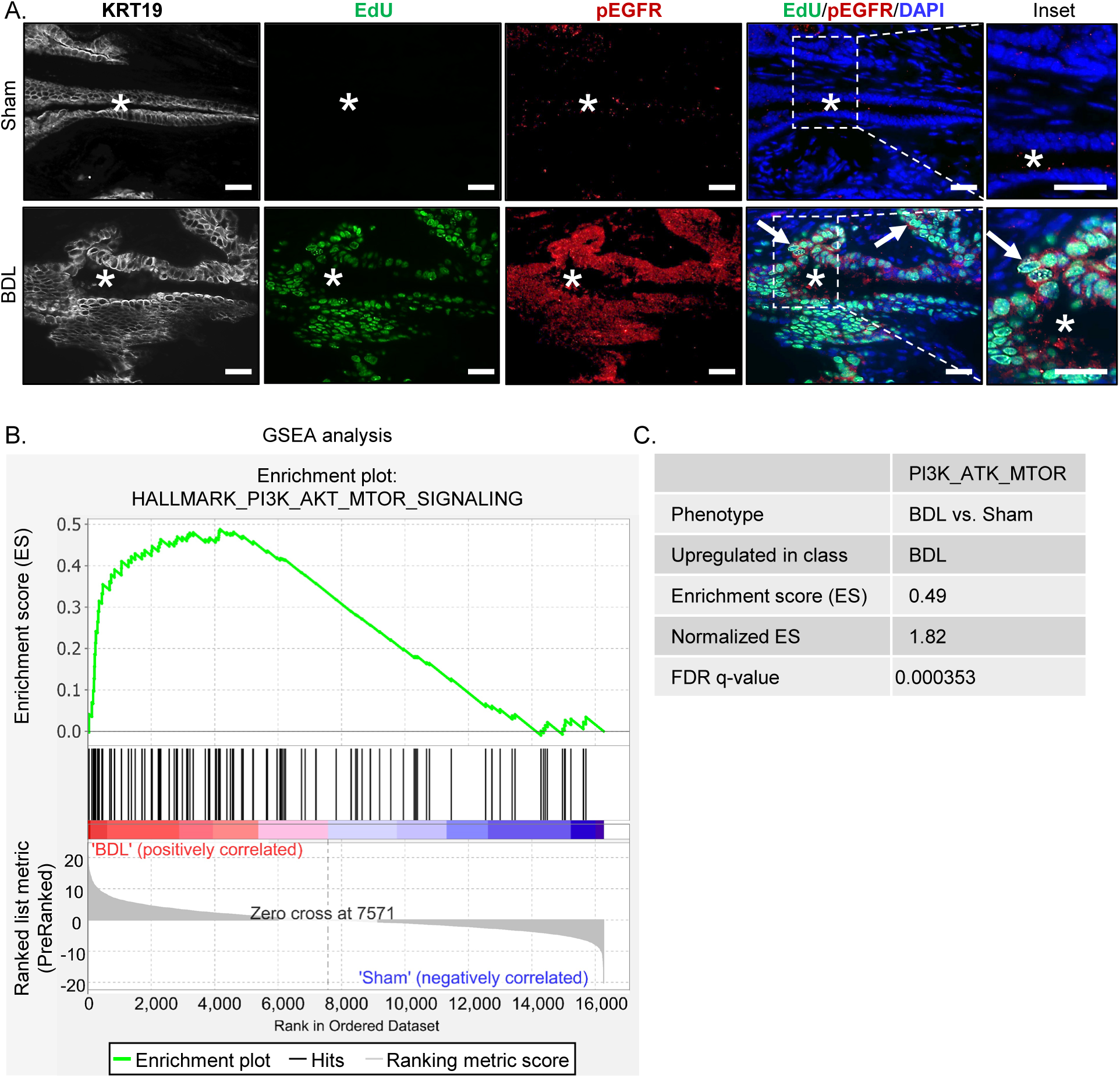
EGFR signaling is activated in the obstructed EHBD. Immunostaining for EGFR activation by Phospho-EGFR (Tyr1068) (pEGFR, red), proliferation (EdU, green), and nuclei (DAPI, blue) 24 hours post-surgery (A). Gene set enrichment analysis (GSEA) using bulk RNA-seq datasets from sham and bile duct ligated (BDL) mouse EHBDs for PI3K/AKT/MTOR signaling (B) and enrichment scores from GSEA analysis (C). Asterisks mark EHBD lumen. Arrows mark pEGFR/EdU co-positive cells. Scale bars, 25μm.

### EGFR-binding ligands promote biliary proliferation in EHBD organoids

To determine the effects of EGFR-binding ligands on EHBD cholangiocytes, we used mouse and human EHBD organoids (6, 29) after confirming that both mouse and human biliary organoids express the genes encoding EGF signaling components including *EGFR* (**Suppl. Fig. 2**). We treated mouse organoids with recombinant EGFR ligands that were upregulated post-BDL (**Fig. 3A**) and observed a dose-dependent growth response to recombinant TGFα, BTC, HB-EGF and EREG (**Fig. 3B-D**). These pro-proliferative responses to EGFR ligands were confirmed in three different mouse and human biliary organoid lines (**Suppl. Fig. 3 and Fig. 4**). We also observed significant increases in organoid establishment rates (organoid number) from BTC and EREG ligands in mice and all ligands in humans, suggesting that these ligands support progenitor cell function (**Suppl. Fig. 3C, E** and **Fig. 4B-F**). Upon treatment with different EGFR ligands, organoids maintained their standard cystic phenotype. Notably, the pro-proliferative effect required 10-100-fold lower concentrations of TGFA, BTC, and HB-EGF (**Suppl. Fig. 3B-D**) as compared with EREG and AREG (**Fig. 3E-F**), with the later showing minimal effect even at the higher dose in mouse organoids. This was consistent with prior work showing different affinities of EGFR ligands (12). We next tested effects of NRG1, the ligand for ERBB3, as it was the only neuregulin significantly increased post-BDL, and observed increased organoid growth at high NRG1 concentrations (**Suppl. Fig. 4**). Thus, our data suggested the interchangeable role of EGF ligands upregulated post-BDL in the promotion of biliary proliferation in organoids.

**Fig. 3.**
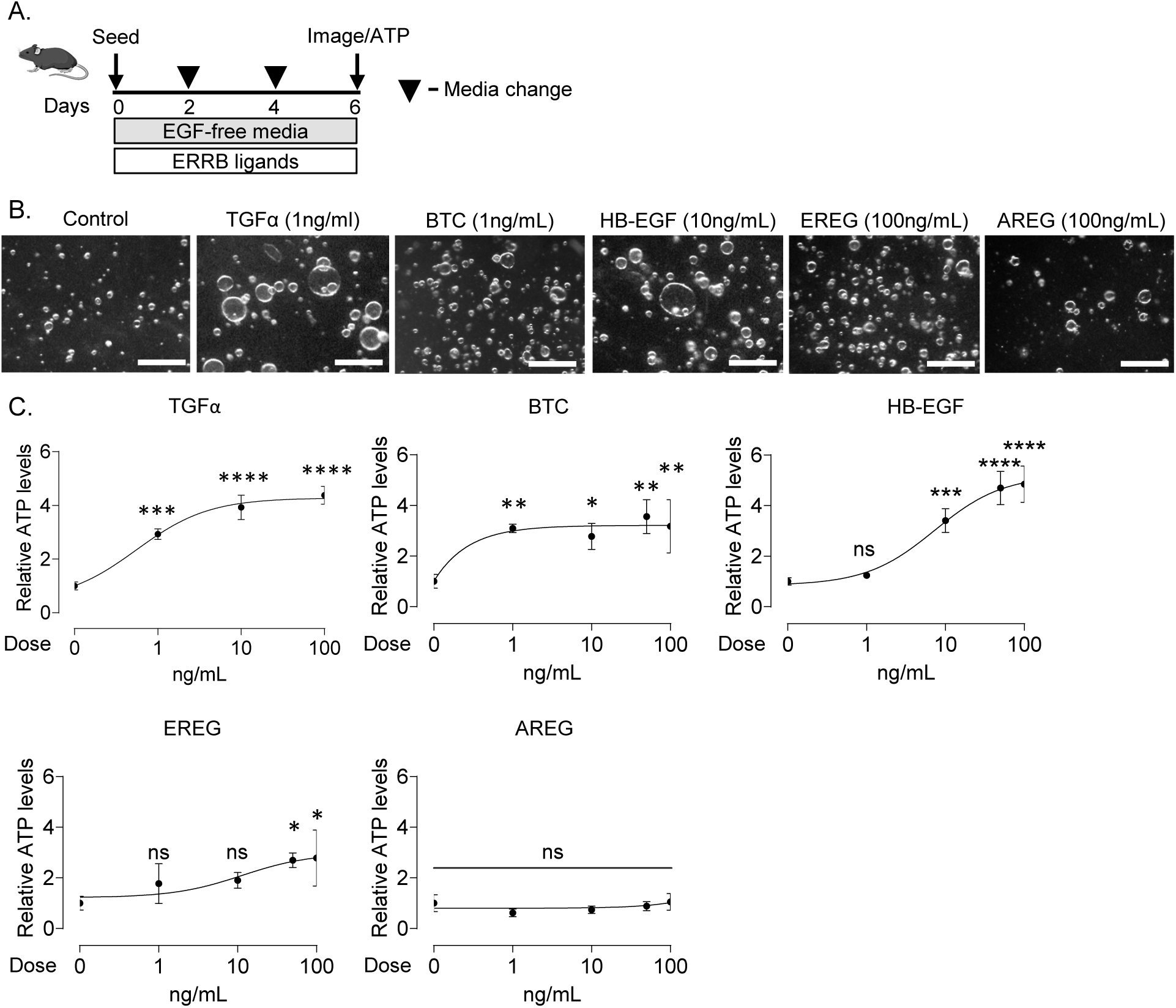
Enhanced growth of mouse EHBD organoids in the dose-dependent response to EGF signaling ligands. Experimental schematic for organoids treated with various epidermal growth factor (EGF) signaling ligands (A). Representative images from control or EGF signaling ligand-treated mouse extrahepatic bile duct (EHBD) organoids. Mouse EHBD organoid growth (ATP measurement) was examined in response to a range of doses for EGF signaling ligands TGFα, BTC, HB-EGF, EREG and AREG (C) n = 3-4 technical replicates. A one-way ANOVA with Dunnett’s multiple comparisons test. The data are presented as the mean ± SD. \**P* < 0.05, \*\**P* < 0.01, \*\*\**P* < 0.001, \*\*\*\**P* < 0.0001, ns – not significant. Scale bars, 500μm.

**Fig. 4.**
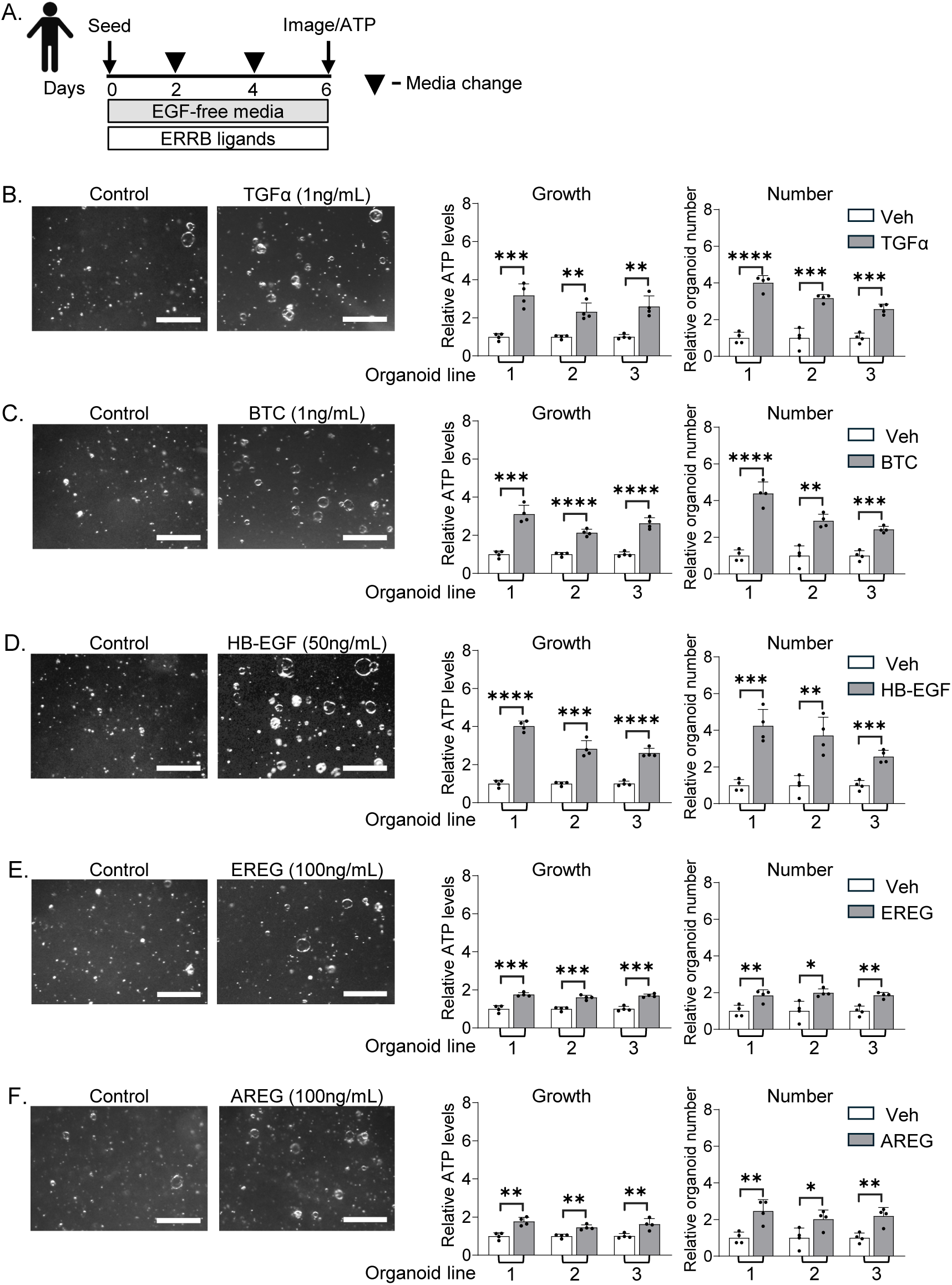
Increased growth and progenitor cell function in human EHBD organoids in response to EGF family signaling ligands. Experimental schematic for organoids treated with various epidermal growth factor (EGF) family signaling ligands (A). Human extrahepatic bile duct (EHBD) organoid growth (ATP measurement) and establishment rate (number) were examined in response to recombinant TGFa (1ng/mL, B), BTC (1ng/mL, C), HB-EGF (50ng/mL, D), EREG (100ng/mL, E), and AREG (100ng/mL, F) ligands. n = 3 biological replicates. Unpaired Student *t*-test. The data are presented as the mean ± SD. \**P* < 0.05, \*\**P* < 0.01, \*\*\**P* < 0.001, \*\*\*\**P* < 0.0001, ns – not significant. Scale bars, 500μm.

### Inhibition of EGFR and not ERBB2 abrogates EHBD biliary proliferation in biliary organoids

TGFα, BTC, HB-EGF, AREG and EREG bind to EGFR extracellular domain and can transmit downstream signals either through EGFR homodimers or through heterodimerization with other EGF family receptors (30). NRG1 binds to ERBB3 and ERBB4. However, ERBB3 lacks kinase activity and relies on heterodimerization with other EGF network receptors for downstream signaling (30). As ERBB4 abundance in EHBDs and cholangiocytes was almost undetectable (**Fig. 1F** and **Suppl. Fig. 2B, D**), we focused on the contribution of EGFR and ERBB2 to the regulation of biliary proliferation by testing an inhibitory paradigm in organoids. EHBD organoids actively proliferate at baseline, unlike quiescent *in vivo* cholangiocytes at homeostasis (6, 25), and both mouse and human biliary organoids expressed *Tgfα, Btc, Hbegf, Areg*, and *Ereg* ligands, and *Egfr* and *Erbb2* receptors (**Suppl. Fig. 2 A, B**). Thus, we hypothesized, that cholangiocytes in organoids maintain their proliferative state through autonomous activation of EGFR and/or ERBB2 signaling. We treated organoids with erlotinib, a Food and Drug Administration (FDA) approved small molecule antagonist of EGFR phosphorylation, and observed decreased growth in mouse and human biliary organoids with an IC^50^ of 0.073 and 0.5 µM, respectively (**Fig. 5 A-E**). Erlotinib also decreased organoid numbers in mouse organoids (**Fig. 5B, C**). These findings further confirmed the direct involvement of EGFR signaling in the promotion of cholangiocyte proliferation.

**Fig. 5.**
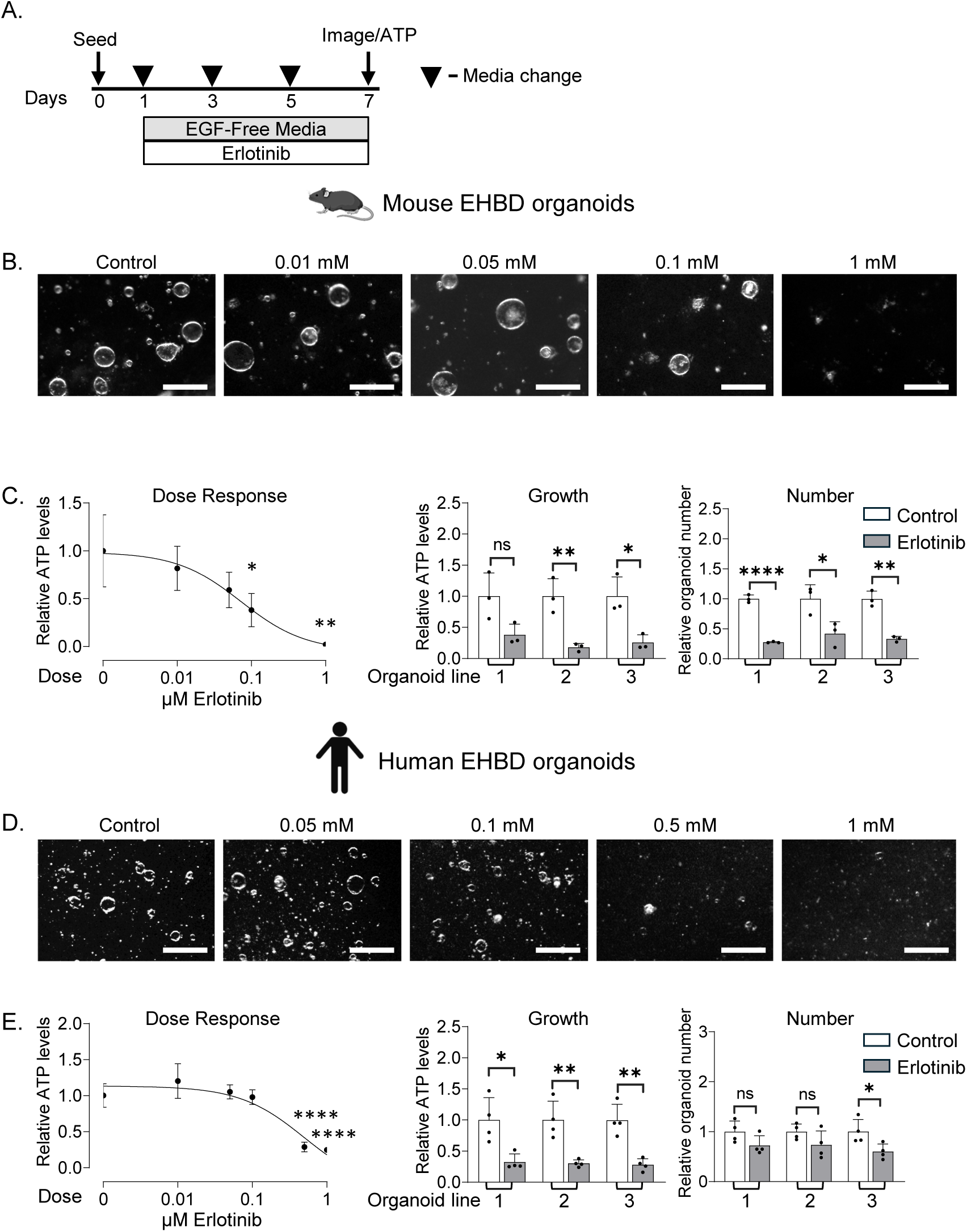
EGFR inhibition attenuates mouse and human EHBD organoid growth. Experimental schematic for organoids treated with the epidermal growth factor receptor (EGFR) inhibitor erlotinib (A). Mouse extrahepatic bile duct (EHBD) organoid growth (ATP measurement) was examined in response to a range of erlotinib doses (B and C, left; n = 3 technical replicates). Mouse EHBD organoid growth and number in response to erlotinib (0.1µM) in 3 biological replicates (C, middle and right). Images and dose response curve of human EHBD organoids treated with a range of Erlotinib concentrations (D and E, left). Growth and number of human EHBD organoids in response to erlotinib (0.5µM) in 3 biological replicates (E). Unpaired Student *t*-test. The data are presented as the mean ± SD. \**P* < 0.05, \*\**P* < 0.01, \*\*\*\**P* < 0.0001, ns – not significant. Scale bars, 500μm.

ERBB2 lacks an extracellular ligand-binding domain, however it can be activated by EGFR ligands through dimerization with EGFR (30). Therefore, to evaluate the potential contribution of ERBB2 to biliary proliferation we used trastuzumab, an FDA approved recombinant humanized anti-ERBB2 antibody (31, 32) (**Fig. 6A**). We utilized human organoids for this experiment as trastuzumab does not target mouse cells. Trastuzumab had no effect on human biliary organoid growth or cell number (**Fig. 6 B-C**), while it effectively inhibited the growth of ERBB2-overexpressing breast cancer cells (23), which were used as a positive control (**Suppl. Fig. 5**). This suggested that ERBB2 is not required for biliary organoid proliferation.

**Fig. 6.**
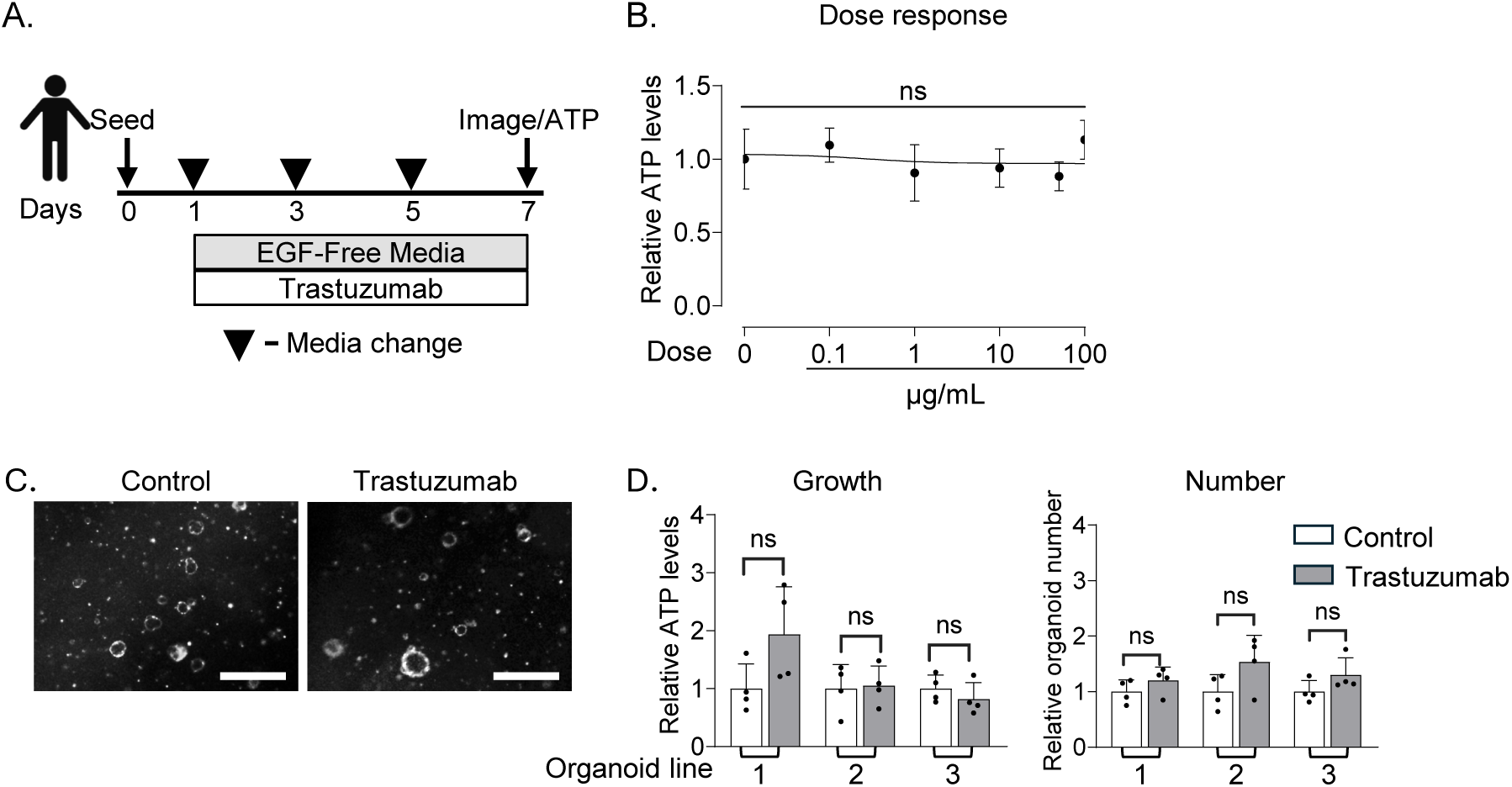
ERBB2 inhibition does not alter human EHBD organoid growth. Experimental schematic for human organoids treated with the ERBB2 inhibitor trastuzumab (A). Dose response curve for trastuzumab treatment concentrations from 0.1μg/mL - 100μg/mL (B). Human extrahepatic bile duct (EHBD) organoid images, growth, and number after treatment with either control (isotype) or trastuzumab (50ug/mL, C). Unpaired Student *t*-test. The data are presented as the mean ± SD. ns – not significant. Scale bars, 500μm.

### Inhibition of EGFR activation in mice results in the dampened biliary proliferative response in obstructed EHBDs

We demonstrated upregulation of EGFR signaling in ligated mouse EHBDs in association with biliary hyperplasia (**Fig. 1, 2**). We also showed that EGFR signaling promotes cholangiocyte proliferation *in vitro* (**Fig. 5**). To determine if EGFR activation is required for the biliary proliferative response *in vivo*, we treated mice undergoing BDL with the EGFR inhibitor erlotinib (**Fig. 7A, B**). Erlotinib-treated mice that underwent BDL exhibited a significant decrease in the number of proliferating (EdU+) cholangiocytes (KRT19+) as compared with vehicle-treated BDL mice (**Fig.7 C, D**). Thus, we demonstrated that EGFR signaling promotes EHBD regeneration through support of biliary proliferation after BDL-induced EHBD injury.

**Fig. 7.**
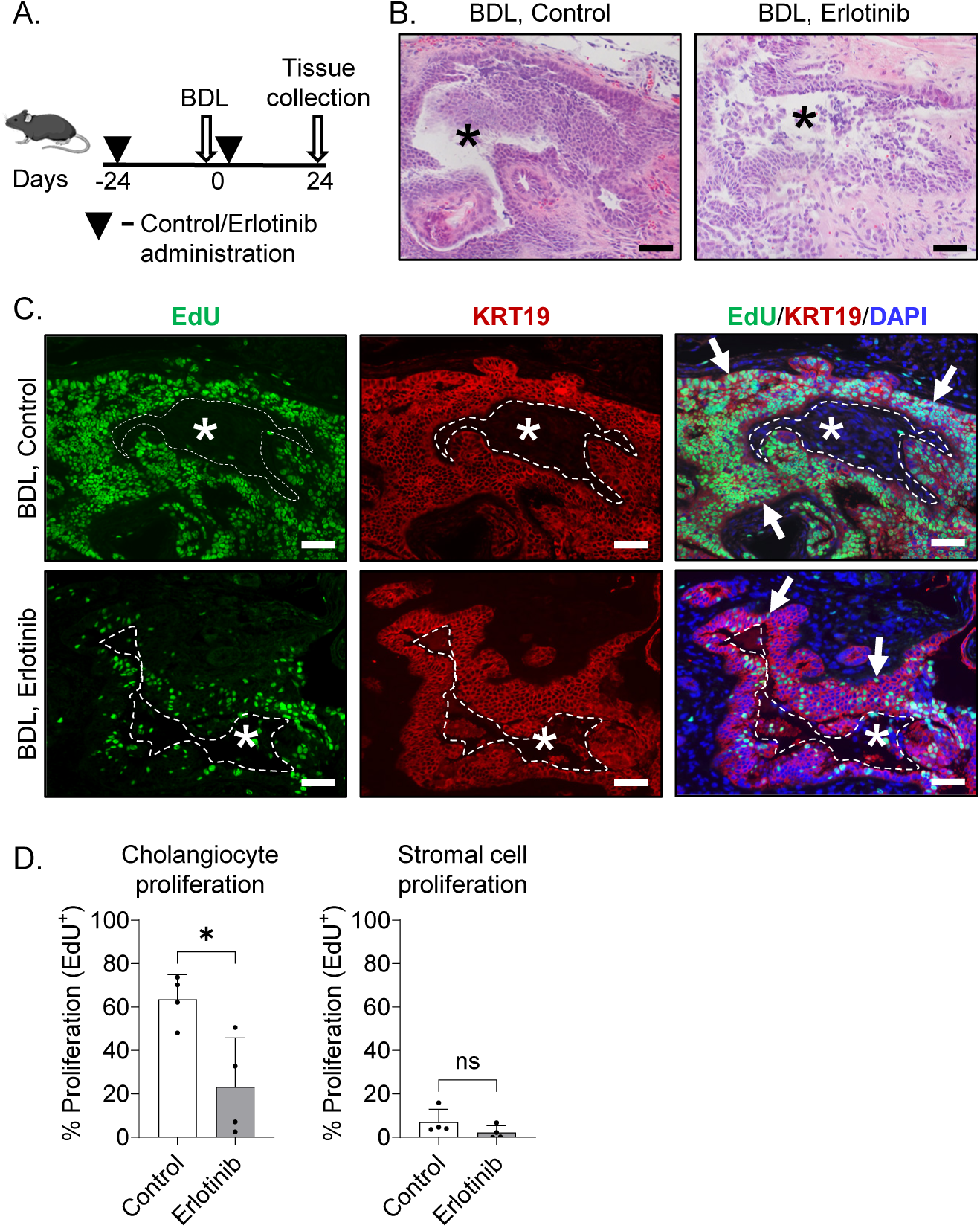
EGFR inhibition decreases cholangiocyte proliferation in injured mouse EHBDs. Experimental schematic for mice treated with vehicle (control) or epidermal growth factor receptor (EGFR) inhibitor erlotinib and undergoing bile duct ligation (BDL) surgery (A). H&E images of mouse extrahepatic bile ducts (EHBDs) 24 hours post-BDL surgery (B). Mouse BDL sections examined with immunofluorescence for proliferation (EdU, green), keratin 19 (KRT19; cholangiocytes, red), and nuclei (DAPI, blue) (C). Morphometric analysis of proliferation in the epithelial (KRT19+ cholangiocyte) and stromal compartments (D). n = 4 mice/group. Unpaired Student *t*-test. The data are presented as the mean ± SD. \**P* < 0.05, ns – not significant. Asterisks mark EHBD lumen. Arrows mark proliferating cholangiocytes. Scale bars, 50μm.

## DISCUSSION

This study investigated the role of EGFR signaling in the cholangiocyte proliferative response to obstructive EHBD injury. While there is a body of work focused on the regulation of epithelial cell responses to injury in the liver and intrahepatic bile ducts, mechanisms regulating EHBD responses to damage are poorly described.

Understanding the regulation of cholangiocyte proliferation can inform approaches to promote biliary regeneration and unveil potential mechanisms of uncontrolled proliferation during biliary carcinogenesis. Our study implicated EGFR signaling in EHBD regeneration after bile duct obstruction by contributing to the biliary proliferative response (**Fig. 8**). This effect involves upregulation of EGFR ligand abundance and phosphorylation of the EGFR receptor in cholangiocytes with enrichment of the downstream signaling pathways, especially along the PI3K/AKT/MTOR axis.

**Fig. 8.**
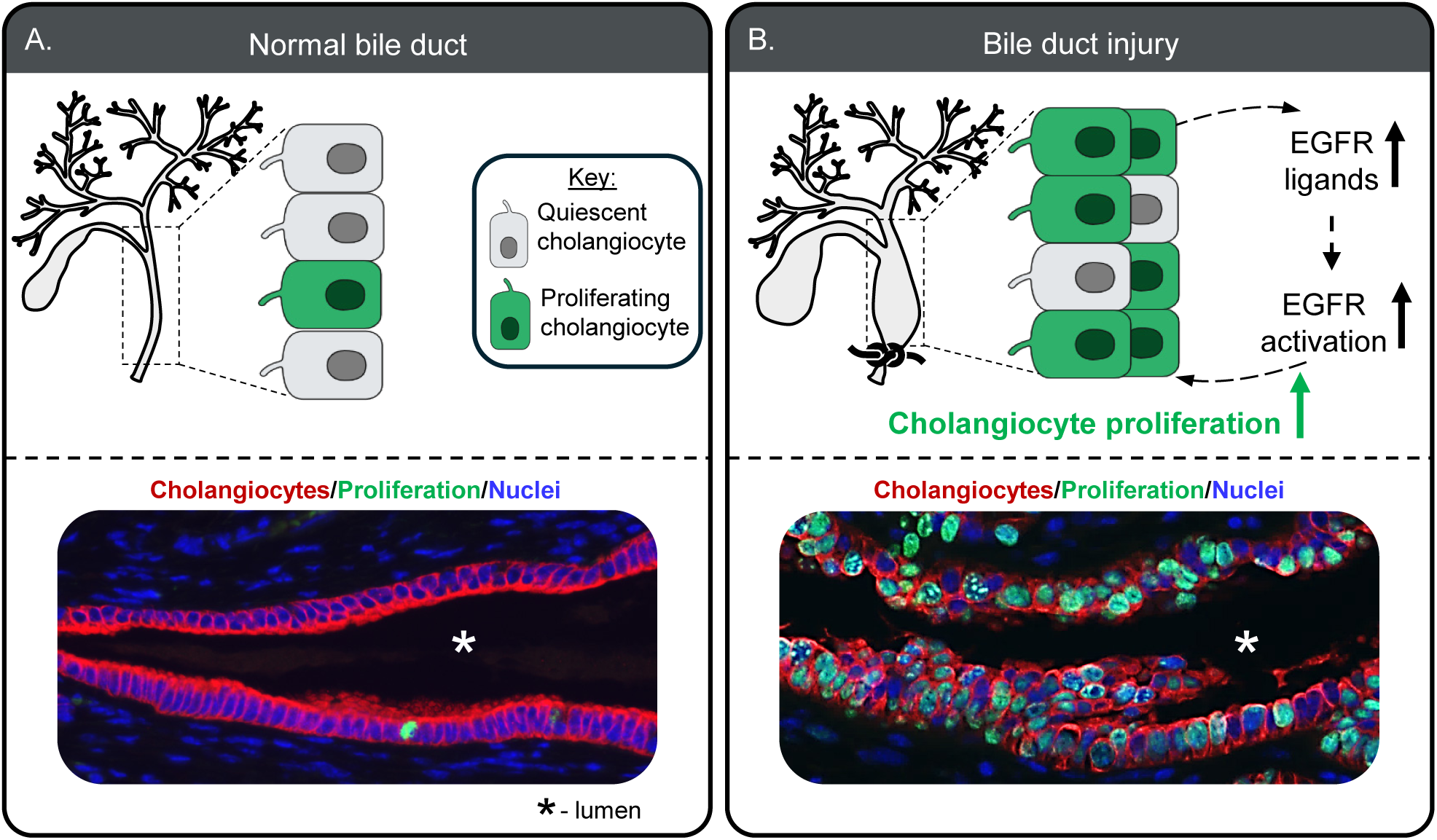
The proposed model of EGFR signaling contribution to EHBD regeneration with biliary proliferation after injury. Cholangiocytes are quiescent in the extrahepatic bile duct (EHBD) at homeostasis (A). EHBD injury with obstruction results in upregulation of EGFR ligands and EGFR activation, which promotes cholangiocyte proliferation (B).

The activation of EGFR by ligands can elicit distinct ligand, organ system, and pathologic condition-dependent responses (33). In the gut, EREG was shown to promote intestinal organoids with striking villus-like structures, while exposure to EGF was shown to stimulate formation of progenitor-like cystic organoids (20). In the liver, EGF directs bile duct morphogenesis and branching in bioengineered bile ducts (34), while AREG inhibits bile acid-induced hepatocyte apoptosis, protects against BDL-induced liver cholestasis, and induces hepatocyte mitosis after partial hepatectomy in mice (15, 35). TGFα, and not EGF or HB-EGF, was suggested to mediate EGFR activation promoting CCA cell proliferation (36). These studies demonstrate the diverse roles EGFR ligands play in tissue maintenance and disease processes throughout the digestive system.

In the EHBD, we showed that all EGFR ligands that were overexpressed in obstructed EHBDs promoted cholangiocyte proliferation in human and mouse EHBD organoids, and organoids maintained a cystic structure without any ligand-dependent phenotypic changes. In our experiments we also observed that traditional high-affinity ligands (37) EGF, TGFa, BTC, and HB-EGF induced cholangiocyte proliferation at lower concentrations than low affinity ligands, EREG and AREG. However, BTC and EREG, but not TGFα and HB-EGF, also increased organoid establishment efficiency in mouse organoids suggesting potential BTC and EREG roles in mouse biliary progenitor cell maintenance. Human organoids mounted more prominent responses to EGFR ligand, AREG, as compared with mouse organoids in our studies suggesting species-specific sensitivity of cholangiocytes to EGFR ligands. Interestingly, although *Areg* expression increased substantially post-BDL, recombinant AREG had limited effects on mouse organoid growth, indicating that it might regulate other regenerative responses than cholangiocyte proliferation in mice and have species-specific effects. Surprisingly, the abundance of *Egf*, the prototypic EGFR ligand implicated in liver diseases and often used for organoid media, was relatively low and mapped to EHBD fibroblasts and immune cells at homeostasis, and *Egf* abundance significantly decreased in obstructed EHBDs. This suggested that EGF may not be important for responses to acute EHBD injury but instead may play a more important role during homeostasis (34). This study provides the basis for further exploration into the roles of specific EGFR ligands in cholangiocyte progenitor cell function and differentiation during homeostasis and in other pathological conditions, including more chronic injury.

Receptor tyrosine kinases, which include EGF receptors, have a critical role in cell proliferation, differentiation, and migration. Increased cholangiocyte EGFR phosphorylation was reported in patients with primary sclerosing cholangitis (38). However, EGF family signaling upregulation is more often associated with cancers including bile duct cancer, CCA (39–42), which is characterized by cellular hyperproliferation (8). Thus, studies reported *EGFR* overexpression in 8.1% of biliary tract cancers, *ERBB2* amplification in 17.2% of perihilar CCA and overexpression of both *EGFR* and *ERBB2* in 7% of biliary cancers (42). *ERBB2* amplifications were reported to be more common in extrahepatic CCA as compared with intrahepatic CCA (17.4% and 4.8% respectively) further highlighting the difference in intrahepatic and extrahepatic bile duct biology (43).

The current study showed that inhibition of EGFR with erlotinib decreased the cholangiocyte proliferative response to EHBD injury. Erlotinib also inhibited the growth of human and mouse biliary organoids further suggesting that EGFR activation is important for cholangiocyte proliferation. We considered that ligands activating EGFR could potentially signal through EGFR/ERBB2 heterodimers as ERBB2 lacks a ligand-binding extracellular subdomain and needs to pair with EGFR and ERBB3 for ligand-mediated signals (44). While ERBB3 depends on binding with other EGF family receptors for kinase activity and could potentially bind with ERBB2. As treatment with ERBB2 inhibitor trastuzumab (45) had no effect on biliary organoid proliferation, we concluded that it is likely EGFR, and not ERBB2 or ERBB3, that conducts pro-proliferative signals in EHBD cholangiocytes. Interestingly, a recent Phase III clinical trial suggested survival benefits from the addition of erlotinib to a combination of gemcitabine and oxaliplatin, for patients with CCA (46). In contrast, the inhibition of ERBB2 in CCA yielded disappointing results (47, 48).

In summary, our comprehensive set of experiments using transcriptomic and functional analyses of a mouse model of obstructive bile duct injury, along with human and mouse organoid models demonstrated an essential need for EGFR activation in promotion of the biliary proliferative response in injured EHBDs. We also demonstrated that many EGFR ligands are upregulated to support this regenerative response. By defining the status and landscape of EGF family signaling in the healthy and diseased EHBD, we provide the foundational framework for further studies focused on the EHBD.

## AUTHOR CONTRIBUTIONS

NR and ANC devised the study design, main conceptual ideas, and project outline. ANC, TS, MP, and NR performed experiments. ANC and NR processed and analyzed the data with TS, JT and MP assistance. TF procured human samples. ANC and NR wrote the manuscript.

## ABBREVIATIONS

EGF: epidermal growth factor
EGF receptor: (EGFR)
ERBB2-4: erythroblastoma oncogene B 2-4
CCA: cholangiocarcinoma
EHBD: extrahepatic bile duct
BDL: bile duct ligation
HER1-4: human epidermal growth factor receptors
TGFα: transforming growth factor-α
BTC: betacellulin
HB-EGF: heparin-binding EGF-like growth factor
AREG: amphiregulin
EREG: epiregulin
NRG1-4: neuregulin 1-4
EdU: 5-ethynyl-2′-deoxyuridine
H&E: hematoxylin and eosin
ATP: adenosine triphosphate
KRT19: keratin 19
DAPI: 4′,6-diamidino-2-phenylindole
RNA sequencing: RNA-seq
single-cell RNA-seq: scRNA-seq
GSEA: gene set enrichment analysis
ANOVA: one-way analysis of variance.

## ACKNOWLEDGEMENTS

We want to thank Drs. Linda Samuelson and Benjamin Allen, and Theresa Keeley, M.S., for the critical feedback on this manuscript and Dr. Carole Parent for providing the breast cancer cell line for this study. Schematics were created in https://BioRender.com

