## Supplemental Data for "EGFR activation in cholangiocytes promotes extrahepatic bile duct regeneration after injury"

A. GSEA analysis

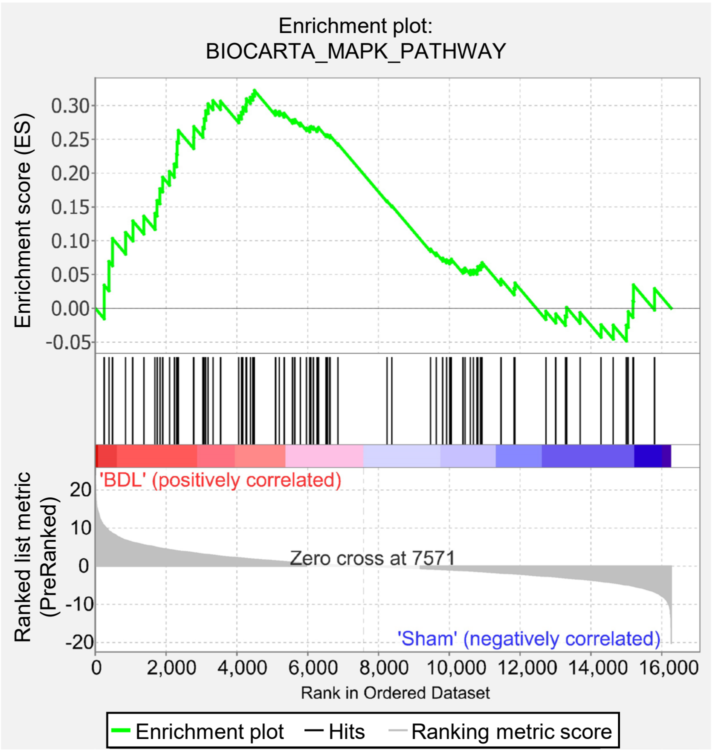


B.

| | MAPK |
|---|---|
| Phenotype | BDL vs. Sham |
| Upregulated in class | BDL |
| Enrichment score (ES) | 0.32 |
| Normalized ES | 1.18 |
| FDR q-value | 0.545 |

**Suppl. Fig. 1: MAPK pathway is not significantly enriched following BDL.** Gene set enrichment analysis (GSEA) using bulk RNA-seq datasets from sham and bile duct ligated (BDL mouse extrahepatic bile ducts (EHBDs) for the MAPK pathway (A) and enrichment scores from GSEA analysis (B).

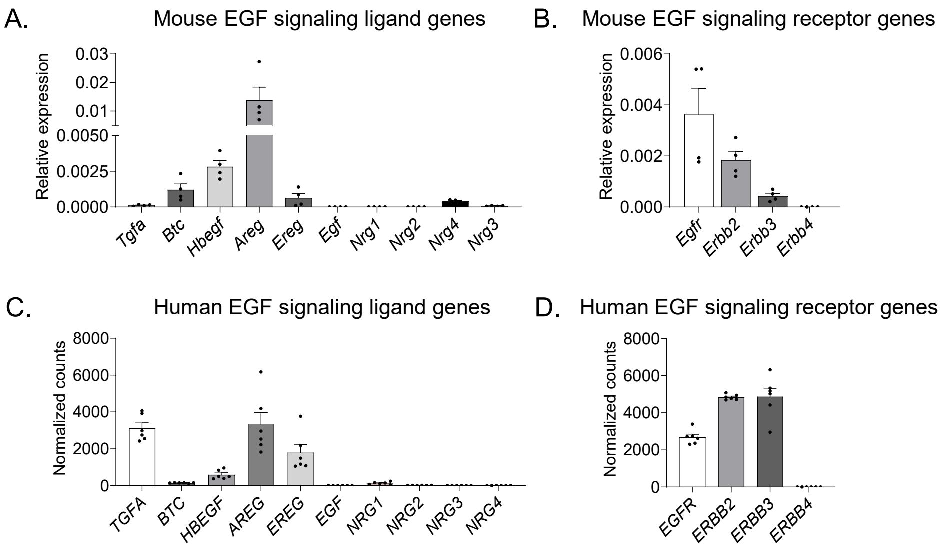


**Suppl. Fig. 2: Genes encoding EGF signaling receptors and ligands are expressed in mouse and human EHBD organoids.** qRT-PCR for epidermal growth factor (EGF) signaling ligands (A) and receptors (B) in mouse extrahepatic bile duct (EHBD) organoids. Analysis of public dataset (E-MTAB-7569) for EGF family ligands (C) and receptors (D) in human EHBD organoids.

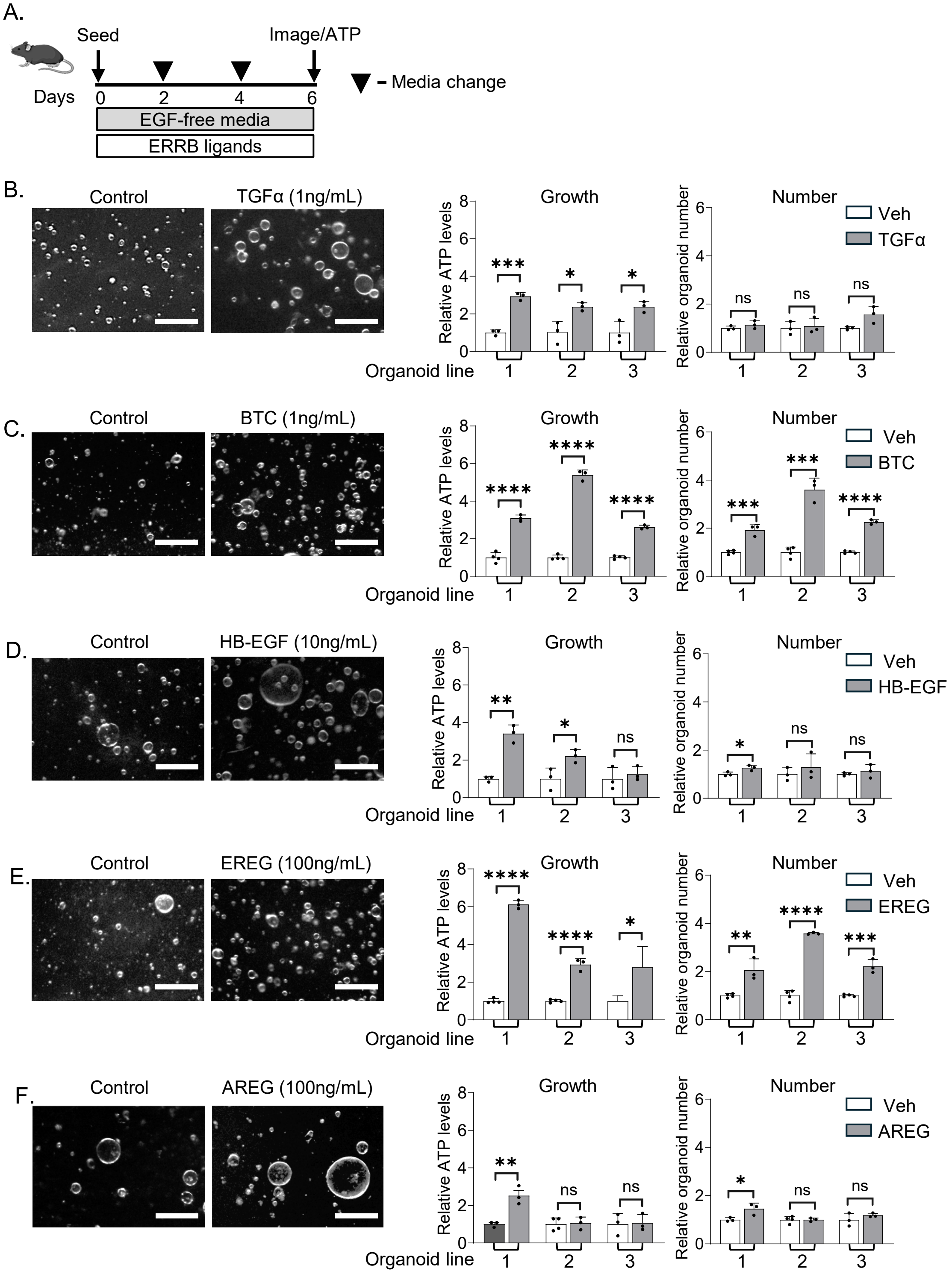


**Suppl. Fig. 3. EGF signaling ligands induce growth and progenitor cell function in mouse EHBD-derived organoids.** Experimental schematic for organoids treated with various epidermal growth factor (EGF) family signaling ligands (A). Mouse extrahepatic bile duct (EHBD) organoid growth (ATP measurement) and establishment rate (number) were examined in response to recombinant TGFα (1ng/mL, B), BTC (1ng/mL, C), HB-EGF (10ng/mL, D), EREG (100ng/mL, E), and AREG (100ng/mL, F) ligands. n = 3 biological replicates. Unpaired Student *t*-test. The data are presented as the mean ± SD. *$P < 0.05$, **$P < 0.01$, ***$P < 0.001$, ****$P < 0.0001$, ns – not significant. Scale bars, 500μm.

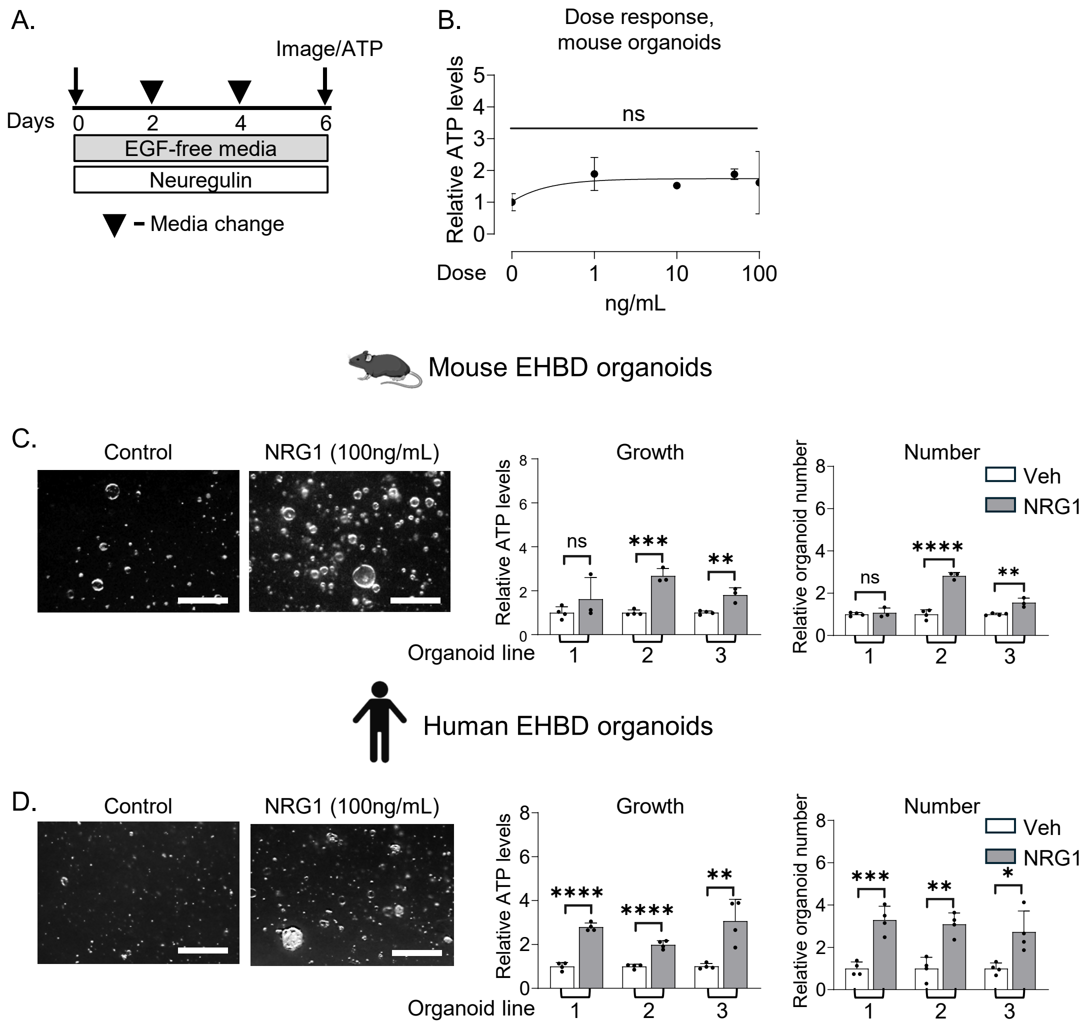


**Suppl. Fig. 4. Increased growth of mouse and human EHBD organoids with the ERBB3 ligand NRG1.** Experimental schematic for organoids treated with the ERBB3 ligand, NRG1 (A). Dose response for NRG1 in mouse extrahepatic bile duct (EHBD) organoids (B). Images, organoid growth, and establishment rate (number) were examined in mouse (C) and human (D) EHBD organoids treated with recombinant NRG1 (100ng/mL). n = 3-4 technical (B, dose-response curve in mouse organoids) and n = 3-4 biological (C and D) replicates. One-way ANOVA with Dunnett's Multiple Comparisons test (B) and Unpaired Student *t*-test (C,D). The data are presented as the mean ± SD. **P* < 0.05, ***P* < 0.01, ****P* < 0.001, *****P* < 0.0001, ns – not significant. Scale bars, 500µm.

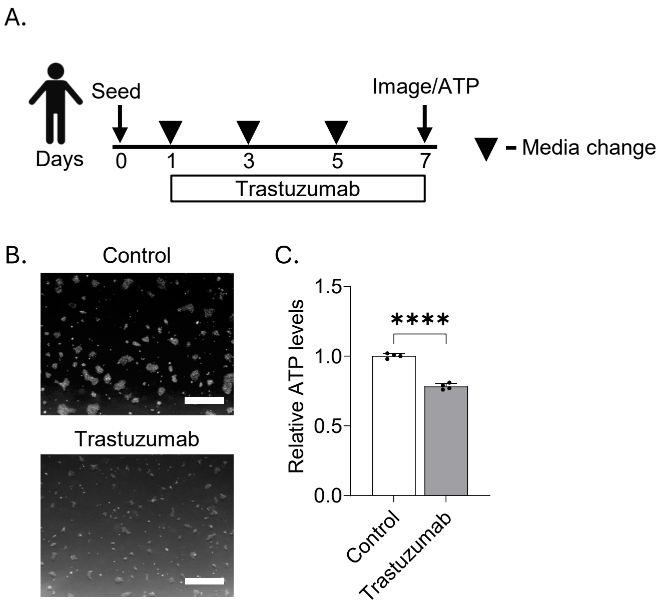


**Suppl. Fig. 5. Trastuzumab inhibits growth in ERBB2 overexpressing breast cancer cells.** Experimental schematic for treatment of ERBB2 overexpressing breast cancer cell line, BT474, with ERBB2 inhibitor trastuzumab. Images (B) and growth (C) in cells treated with control (isotype) or trastuzumab (10ug/mL). Unpaired Student *t*-test. The data are presented as the mean ± SD. ns – not significant. Scale bars, 500μm.

| **Antibody:** | **Host:** | **Company:** | **Catalog#:** | **Concentration:** |
|---|---|---|---|---|
| **Primary:** | | | | |
| KRT19 | Rat | DSHB | Troma III | 1:100 |
| Phospho-EGFR (Tyr1068) | Rabbit | Invitrogen | 44-788G | 1:50 |
| **Secondary:** | | | | |
| Alexa Fluor 555 | Goat | Invitrogen | A21428 | 1:1000 |
| Alexa Fluor 555 | Goat | Invitrogen | A21434 | 1:1000 |
| Biotinylated | Donkey | Vector Laboratories | BA-9400 | 1:8000 |

**Suppl. Table 1. Primary and secondary antibodies.**

| **Gene target:** | **Forward Primer:** | **Reverse Primer:** |
|---|---|---|
| *18s* | 5’- GTAACCCGTTGAACCCCATT -3’ | 5'- CCATCCAATCGGTAGTAGCG -3' |
| *Egfr* | 5'- GCCATCTGGGCCAAAGATACC -3' | 5'- GTCTTCGCATGAATAGGCCAAT -3' |
| *Erbb2* | 5'- CAGCTCGGAGACCTGCTATG -3' | 5'- GTTCGTCCAGGTCCACACAT -3' |
| *Erbb3* | 5'- TCGCCTGGATGTCCTCCTAA -3' | 5'- GGTCACACTCAGCCCGTTTA -3' |
| *Erbb4* | 5'- CAGATCAGGATCGGGAGTGC -3' | 5'- TGGTAAAGTGGAATGGCCCG -3' |
| *Tgfα* | 5’- GCACCCTGCGCTCGGAAGAT -3’ | 5’- TCTGGGATCTTCAGACCACT -3’ |
| *Btc* | 5'- AGCACAGTTGATGGACCCAA -3' | 5'- CAGGAGGGAGTTTGCTCGTC -3' |
| *Hbegf* | 5’- CGGGGAGTGCAGATACCTG -3’ | 5’- TTCTCCACTGGTAGAGTCAGC -3’ |
| *Ereg* | 5'- CAGCACAACCGTGATCCCAT -3' | 5'- CAGACCAGTGTAGCCCACTT -3' |
| *Areg* | 5’- GGGGACTACGACTACTCAGAG -3’ | 5’- TCTTGGGCTTAATCACCTGTTC -3’ |
| *Egf* | 5’- CCTGCCCCCTTCCTAGTTTTC -3’ | 5’- CTCCGTTCTGTTGGTCTACCC -3’ |
| *Nrg1* | 5'- TCTCATCCGAGGCATACACT -3' | 5'- GTCCCAGTCGTGGATGTAGA -3' |
| *Nrg2* | 5'- GCCAGATCCTAAGCAAAAGGC -3' | 5'- GTGGTCTGTAGCTGGCACAT -3' |
| *Nrg3* | 5'- CTATCAAGCACCACAGCCCA -3' | 5'- AGCTGTATAGGCAGGTGGGA -3' |
| *Nrg4* | 5’- GACTGTGGACCATACGACGA -3’ | 5’- GGCCAGTGATGACAGTAGCAG -3’ |

**Suppl. Table 2. Quantitative reverse transcription polymerase chain reaction (qRT-PCR) primers.**
